# Zea mays genotype influences microbial and viral rhizobiome community structure

**DOI:** 10.1101/2023.06.09.544353

**Authors:** Pooja Yadav, Amanda Quattrone, Yuguo Yang, Jacob Owens, Rebecca Kiat, Thirumurugen Kuppusamy, Sabrina E. Russo, Karrie A. Weber

**Author notes:** Corresponding Author: Karrie A. Weber 232 Manter Hall University of Nebraska—Lincoln Lincoln, NE 68588-0118 (402)472-2739. University of Nebraska-Medical Center, Lincoln, NE.

## Abstract

Plant genotype is recognized to contribute to variations in microbial community structure in the rhizosphere, soil adherent to roots. However, the extent to which the viral community varies has remained poorly understood and has the potential to contribute to variation in soil microbial communities. Here we cultivated replicates of two different genotypes of *Zea mays parviglumis* and *Z. mays* genotype B73 in a greenhouse and harvested the rhizobiome (rhizoplane and rhizosphere) to identify the abundance of cells and viruses as well as apply 16S rRNA gene amplicon sequencing and genome resolved metagenomics to identify the rhizobiome microbial and viral community. Our results demonstrate that viruses exceeded microbial abundance in the rhizobiome of *parviglumis* and B73 with a significant variation in both, the microbial and viral community between the two genotypes. Of the viral contigs identified only 4.5% (n =7) of total viral contigs were shared between the two genotypes, demonstrating that plants even at the level of genotype can significantly alter the surrounding soil viral community. An auxiliary metabolic gene associated with glycoside hydrolase (GH5) degradation was identified in one viral metagenome-assembled genome (vMAG) identified in the B73 rhizobiome infecting Propionibacteriaceae (Actinobacteriota) further demonstrating the viral contribution in metabolic potential for carbohydrate degradation and carbon cycling in the rhizosphere. This variation demonstrates the potential of plant genotype to contribute to microbial and viral heterogeneity in soil systems and harbor genes capable of contributing to carbon cycling in the rhizosphere.

## Introduction

Plant roots create a carbon and nutrient-rich environment in soils supporting microbiota in the zone adjacent to the root epidermis and the soil adhering to plant roots termed the rhizoplane and rhizosphere or together referred to as the rhizobiome [1]. It is within this region where a diversity of microbiota including Bacteria, Archaea, Eukarya and viruses are recognized as members of the rhizobiome contributing to below ground processes [2]. The microbial diversity of this environment has the potential to be impacted by viral infection where viral mediated cell lysis is recognized to impact microbial abundance in soils [3]. Viruses are abundant in soils ranging from 10^7^ to 10^9^ viruses g^-1^ of soil [4–6] and virus abundance can often exceed that of microbes [7]. To date a few studies have virus abundance and predation in the rhizosphere [8, 9], but the impact of these processes in this dynamic region in soils remains poorly studied.

Virus infection can result in host cell lysis through a lytic or lysogenic infection thereby, reducing host cell abundance. The lysis of these microbes results in the release of cell lysate which serves as an easily accessible carbon and nutrient source supporting the growth of other microbial members [10]. Hence, viral infections of microbial host cells can influence the metabolic functions of the microbial community. This may occur due to shifts in taxonomic composition and change in metabolic potential caused by differential viral predation or through the acquisition of auxiliary metabolic genes (AMGs) [11–15] resulting from viral infection. AMGs are viral acquired genes from their host, not required in viral replication but allow viruses to directly manipulate host metabolism [16]. Some well-known examples of AMGs acquired by viruses have been inferred to play a role carbon [15], sulfur [17], and nitrogen [18] cycling. Together, these changes in relative abundance of microbial composition and introduction of functional genes like AMGs can alter the metabolic potential of microbial community by metabolically reprogramming their hosts, and/or expression of AMGs [9, 15, 19, 20]. While, the abundance of viruses in the rhizosphere has been reported before [8, 9, 21], the implication of viral impacts on the host abundance and metabolic potential remains understudied. Given the absence of universal marker genes across viruses, metagenomic studies are required to shed light on the role of viruses in rhizobiome and, how viruses impact the abundance of the host cells, and their potential to reprogram host metabolism [22].

In the carbon-rich region surrounding the plant root, the rhizobiome community structure varies between plant species [23] and genotype [24], owing to the influence of the plant, including variation in root exudate profiles [25]. The variation in root exudate profile secreted by host genotype and the resulting substrate-driven or bottom-up shifts in the microbial community structure are based on differential production of root exudates and community utilization [26, 27]. In addition to substrate driven controls on rhizobiome structure, viruses are recognized to infect microorganisms in the rhizosphere [8, 9]. Viruses require metabolically active hosts for replication and thus the patterns of viral communities are linked to active host cells [28–31]. While physiochemical heterogeneity in soils can alter viral abundance where it is recognized that soil moisture and pH can influence viral community structure [32–34], variation in plant roots also has the potential to contribute to viral community structure. In soils viruses impact host abundance and metabolic potential by infecting metabolically active organisms driving carbon-cycling [15, 35], and play a role in biogeochemical processes in the rhizosphere [9, 36]. The region surrounding the root is one are where microorganisms are recognized to be metabolically active owing to the production of carbon-rich plant root exudates.

Here we test the hypothesis that the microbial and viral rhizobiome community varies between two *Z. mays* genotypes, *Z. mays ssp. parviglumis* (teosinte) and *Z. mays* genotype B73 (domesticated corn). Seeds of each genotype were sterilized, germinated under axenic conditions and transplanted into a homogenized soil matrix and cultivated to the V4-V8 growth stage in a greenhouse under controlled conditions. Plants were harvested for phenotyping and rhizobiome samples were collected to enumerate cells and viruses as well as identify microbial community members using 16S rRNA gene amplicon sequencing and genome resolved metagenomics from shotgun metagenome sequence data. Variation within the microbial and viral communities between *Z. mays* genotypes was statistically determined. The virus linked microbial hosts in the rhizobiome of the two genotypes rendered differences in host abundance. Assembled and annotated viral contigs were screened for functional genes defined as AMGs and included the potential to degrade carbohydrates. Here we report differential microbial and viral community assemblage in the rhizobiome between two *Z. mays* genotypes, *parviglumis* and B73.

## Materials and methods

### Seed source, germination, and Z. mays growth

Seeds of *Z. mays parviglumis* and B73 were obtained from the University of Nebraska, Lincoln in-house collection [37]. Twenty-four *parviglumis* and B73 seeds were sterilized and germinated in sterilized petri-plates. From among the 24 seedlings, 12 seedlings were transplanted into 1.8 L pots containing homogenized sieved prairie soil mixed with autoclaved agricultural field sandy-soil (60:40; mass/mass; Supplementary Information) [38]. Plants were cultivated in a greenhouse with an average 13-hour day length for 22 days at 31°C with relative humidity between 60.2 - 97.6 %. Every three days plants were watered and fertilized with 25 mL of 25% Hoagland’s solution [39]. Six biological replicates at the same growth stage (V4-V8) were selected for harvesting.

### Rhizobiome harvesting and sample collection

Plant roots and rhizobiome were harvested by gently cutting the pot to minimize soil and root disruption and carefully separated from bulk soil as previously described [37]. The roots were submerged in a known volume of sterile, nucleic acid and enzyme-free 1X phosphate buffered saline (PBS) and sonicated (Branson 450D, 30% amplitude, 0.3 s duty cycle) for one minute, three times at 30 seconds intervals [40, 41] to harvest the rhizobiome, rhizoplane and rhizosphere. The rhizobiome suspension was centrifuged (800 X *g*). A known volume of the supernatant was filtered through 0.45 µm PVDF and 0.2 µm PVDF membrane, and fixed with formaldehyde (0.5 v/v final concentration) for cell and virus enumeration [42, 43]. The remaining suspension was centrifuged (3000 X *g* for 10 minutes) and supernatant was strained through a 40 mm sterile cell strainer (Fisherbrand™ product #22363547) to remove plant cells and root debris. The supernatant was collected on 0.2 µm polycarbonate membranes (product #GTTP04700) through vacuum filtration (≤ 2.5 bar). The rhizosphere pellet was collected by another round of centrifugation at 10,000 X *g* for 20 minutes. The membrane and pellet were flash frozen in liquid nitrogen and stored at -80 °C. Following rhizobiome collection root phenotype characteristics were determined (Supplementary Information).

### Nucleic acid extraction, 16S rRNA gene amplicon, and shotgun metagenomic sequence library preparation

All reagents and materials used in sample processing were treated or prepared in 0.1 % diethylpyrocarbonate or RNase Away. Nucleic acids were extracted from a known mass of the rhizosphere pellet (10% w/w of total rhizosphere and rhizoplane) and the membrane collected cells using a modified Griffith’s method as previously described [43, 44]. DNA was quantified and purity was checked as previously described (Westrop et al. 2023). A quantified mass of DNA extracted from the pellet and the filter (2.5 % of total DNA (ng) were proportionally pooled to represent the total rhizobiome community. The DNA quantity was normalized per gram rhizosphere and sent for sequencing.

The 16S rRNA gene amplicon sequencing library was prepared by the University of Minnesota Genomics Center, Minnesota using V4 and V5 regions of the 16S rRNA gene primers 515F and 806R as described [45, 46]. The samples were paired-end sequenced (2 X 300 bp) using Illumina MiSeq600 cycle v3 kit to a sequencing depth of 72,000 reads per sample. The 16S rRNA gene amplicon sequence data was analyzed using the DADA2 package to obtain amplicon sequence variants (ASVs) as described [47] and calculate diversity measures (Supplementary Information). The ASVs were filtered for spurious sequences as described and was analyzed using Phyloseq [48].

The shotgun metagenome sequence library was prepared using the Nextera XT DNA library kit (Illumina, California, USA) as previously described [43]. The average final library size was 300 bp with an insert of 180 bp and a read length configuration of 150 PE for 40 M PE reads per sample (20 M in each direction). Sequence data were quality checked (Supplementary Information) and clean reads were used to generate de novo co-assemblies using Megahit [49]. Assembled contigs were binned using MetaBat2 (v2.15) [50], MaxBin2 (v2.2) [51], and CONCOCT (v1.1) [52] and dereplicated using DAS_Tool (v1.1) [53]. The bins were quality checked using CheckM (v1.1) [54] and manually curated using Anvi’o (v7.0) [55] as described [43]. The final set of MAGs were taxonomically identified and used to estimate relative abundance as described (Supplementary Information).

Putative viral contigs were identified using VirSorter2 (v2.1) [56] and DeepVirFinder [57] (Supplementary Information). A pairwise comparison with ≥ 95 % average nucleotide identity (ANI) across ≥ 85 % alignment coverage [58, 59] was used to assess genome similarity in the viral community of *parviglumis* and B73. Viral contigs ≥10 Kbp are described as a vMAG [60]. Viral genome coverage was normalized as performed for MAGs and used as a proxy for relative abundance as previously described [15, 58, 61].

Virus-host linkages were established utilizing a combination of three different methods, clustered regularly interspaced short palindromic repeats (CRISPR) sequences [62], genome similarities [63], and tetranucleotide frequencies (threshold distance < 0.001) between host and viral contig (Supplementary Information) [15, 63–66]. Host genome metabolic potential determined using METABOLIC (v4.0) [67], DRAM (v1.2) [68], and FeGenie (v1.0) [69] at the MAG level. Viral contribution in the host metabolic processes was investigated by screening viral encoded AMGs in *parviglumis* and B73 using DRAM-v.

### Statistical analyses

All statistical analyses were conducted in R version 4.1.2 (2021-11-01). Differences between *Z. mays* genotype in bacterial and virus abundances were tested using t-tests assuming unequal variances. The correlation of cells and virus abundances were analyzed using Mantel’s test and linear regression (Supplementary Information). Relationships between the microbial and viral community structure were analyzed using principal coordinate analysis (PCoA) with Bray Curtis dissimilarities as implemented in the vegan package [70]. Differences between *Z. mays* genotypes in the microbial and viral community composition were evaluated using permutational multivariate analysis of variance (perMANOVA) using adonis2.

## Results

### Virus to cell ratio in the parviglumis and B73 rhizobiome

Virus abundance exceeded cell abundance in the rhizobiome collected from both *parviglumis* and B73 (VCR of 2.24 ± 0.21 and 1.52 ± 0.12, respectively) (Fig. 1) and was positively correlated with cell abundance (*parviglumis p* = 0.001; B73 *p* = 0.008). Cell abundance in the *parviglumis* and B73 rhizobiome totaled (1.34 ± 0.30) x 10^7^ cells g^-1^ and (1.54 ± 0.16) x 10^7^ cells g^-1^ respectively but was not statistically different between genotypes (*p >* 0.05, Fig. 1a). Similarly, *parviglumis* and B73 rhizobiome virus abundance of *parviglumis* (2.89 ± 0.55) x 10^7^ g^-1^ and B73 (2.28 ± 0.20) x 10^7^ g^-1^ rhizosphere (Fig. 1b) exceeded cell abundance and was not statistically different between genotypes (*p > 0.05*). Cell and virus abundance did not differ significantly between genotypes, even when standardized by plant phenotypic parameters including root length, surface area, volume, density, fresh weight, or biomass (Supplementary Fig. S1 and Fig. S2). Cell abundance gram^-1^ of rhizobiome collected from *parviglumis* and B73 was positively correlated with the rhizosphere mass. While cell or virus abundance gram^-1^ of rhizobiome collected was not statistically different between genotypes, the VCR significantly varied (*p = 0.02*) between genotypes indicating variation between genotypes.

**Fig. 1.**
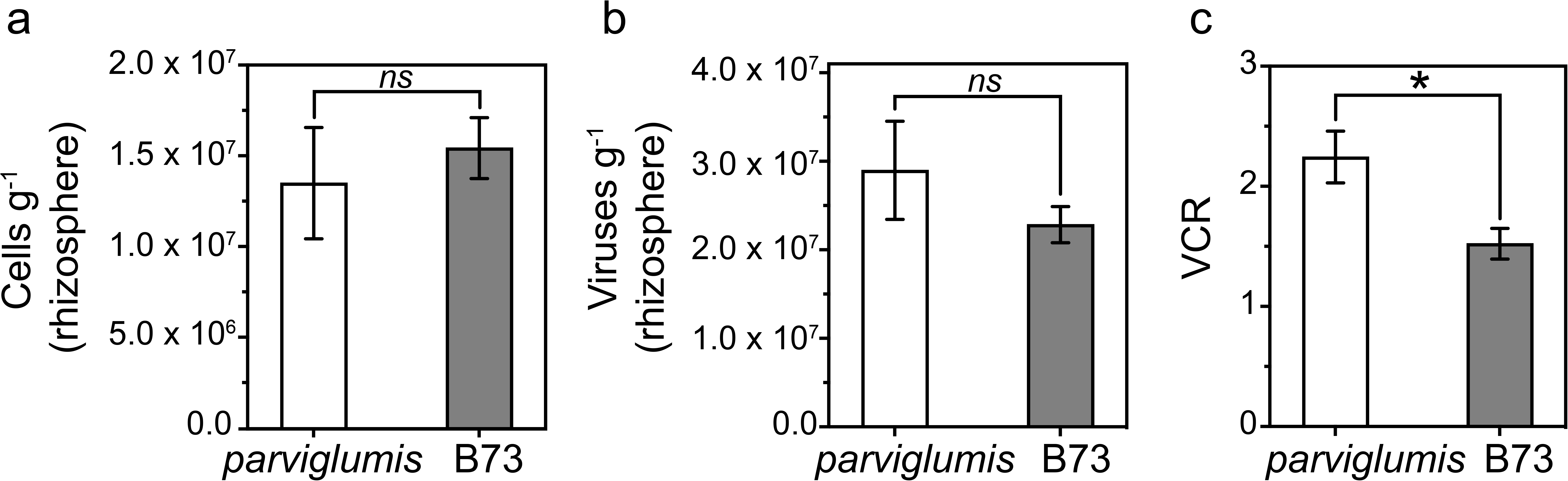
Cell and virus abundance in the parviglumis and B73 rhizobiome. (a) Cell abundance per gram of soil in the rhizoplane and rhizosphere referred to a rhizobiome of biological replicates of *parviglumis* (n=5) and B73 (n=6). (b) Virus abundance per gram of the rhizobiome soil. (c) Virus to cell ratio (VCR) between *parviglumis* and B73. The brackets indicated a non-parametric unpaired t-test with Welch’s correction assuming unequal variances. The asterisk denoted a *p*-value < 0.05 as statistically significant whereas *ns* denoted that the result was not significant. Values reported are the mean of replicates and error bars represent the standard error of measure.

### Microbial community composition of parviglumis and B73 rhizobiome

The rhizobiome microbial community collected from *parviglumis* and B73 revealed 6,317 unique ASVs from clean reads of *parviglumis* (range 34,397 to 96,390) and B73 (range 29,779 to 75,780). The ASVs were broadly classified into 1 archaeal phylum and 17 bacterial phyla (Fig. 2a). The core rhizobiome consisted of Verrucomicrobiota, Proteobacteria, and Actinobacteriota with an abundance in similar proportions accounting for a combined 72.64% and 74.97% of the rhizobiome community in *parviglumis* and B73, respectively. The microbial community identified by 16S rRNA gene amplicon sequencing was statistically different between genotypes. (Fig. 2b; *R^2^* = 0.098, *p* = 0.001) where *Z. mays* genotype accounted for 10.8% of the variation in the microbial community structure (Fig. 2b).

**Fig. 2.**
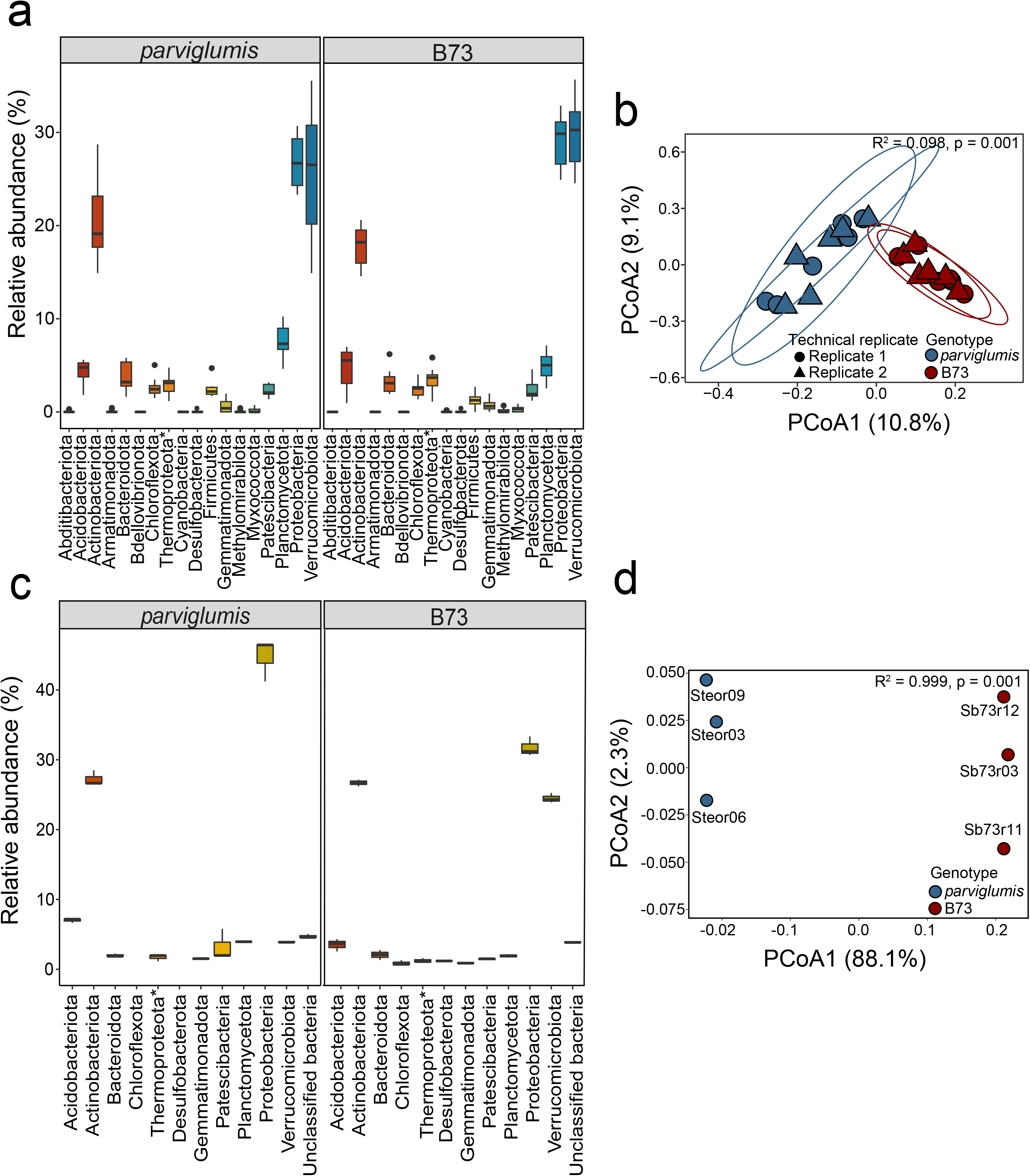
Microbial community composition of Z. mays parviglumis and B73 identified by 16S rRNA gene amplicon sequence variants (ASVs) and metagenome-assembled genomes (MAGs). The microbial community of *parviglumis* and B73 represented (a) a percentage of the relative abundance of the microbial community identified by ASV. (b) PCoA of ASV identified rhizobiome from parviglumis (blue) and B73 (red) in biological and technical replicates with the Bray-Curtis distance matrix. The variance of 56.1% on PCoA1 and 17.4% on PCoA2 (*R^2^* =0.1089, *p*-value 0.047) as determined by perMANOVA. Ellipses denote 95% confidence interval. (c) Relative abundance of MAGs determined using normalized genome coverage (d) PCoA of MAG identified rhizobiome from *parviglumis* (blue) and B73 (red) with Bray-Curtis dissimilarities with 88.1% variance on PCoA1 and 2.3% variance on PCoA2 (*R^2^* =0.999, *p* = 0.001). The box plot represents the interquartile range with whiskers ± 1.5 times of interquartile range and the center depicts the median.

Genome assembly of metagenomic sequence data resulted in MAGs similar to taxa identified at ASV level where reads represented more than 2.5% of the microbial community (Fig. 2c). Biological replicates of the shotgun metagenome sequenced data yielded clean reads in the range of 22.85 to 24.12 million for *parviglumis* and 23.35 to 27.30 million for B73. These data rendered 50 and 61 rhizobiome metagenome-assembled genomes (MAGs) for *parviglumis* and B73, respectively. The *parviglumis* rhizobiome MAGs averaged 42.91 ± 2.49% completeness, totaling 15 medium-quality MAGs and 35 low-quality MAGs and B73 MAGs averaged estimated completeness of 43.60 ± 2.23%, comprising one high-quality, 19 medium-quality, and 41 low quality MAGs. Nine and 11 phyla in the archaeal and bacterial domains were identified in *parviglumis* and B73, respectively (Fig. 3). Similar to 16S rRNA sequencing data, the Thermoproteota was the only archaeal MAG constructed from the rhizobiome of both genotypes. The bacterial MAGs reconstructed from *parviglumis* rhizobiome were Acidobacteriota, Actinobacteriota, Bacteroidota, Desulfobacterota, Gemmatimondadota, Patescibacteria, Planctomycetota, Verrucomicrobiota, and unclassified bacteria. In addition to these taxa, two additional phyla, Chloroflexota and Desulfobacterota, were also reconstructed from B73 rhizobiome (Fig. 2c and 3).

**Fig. 3.**
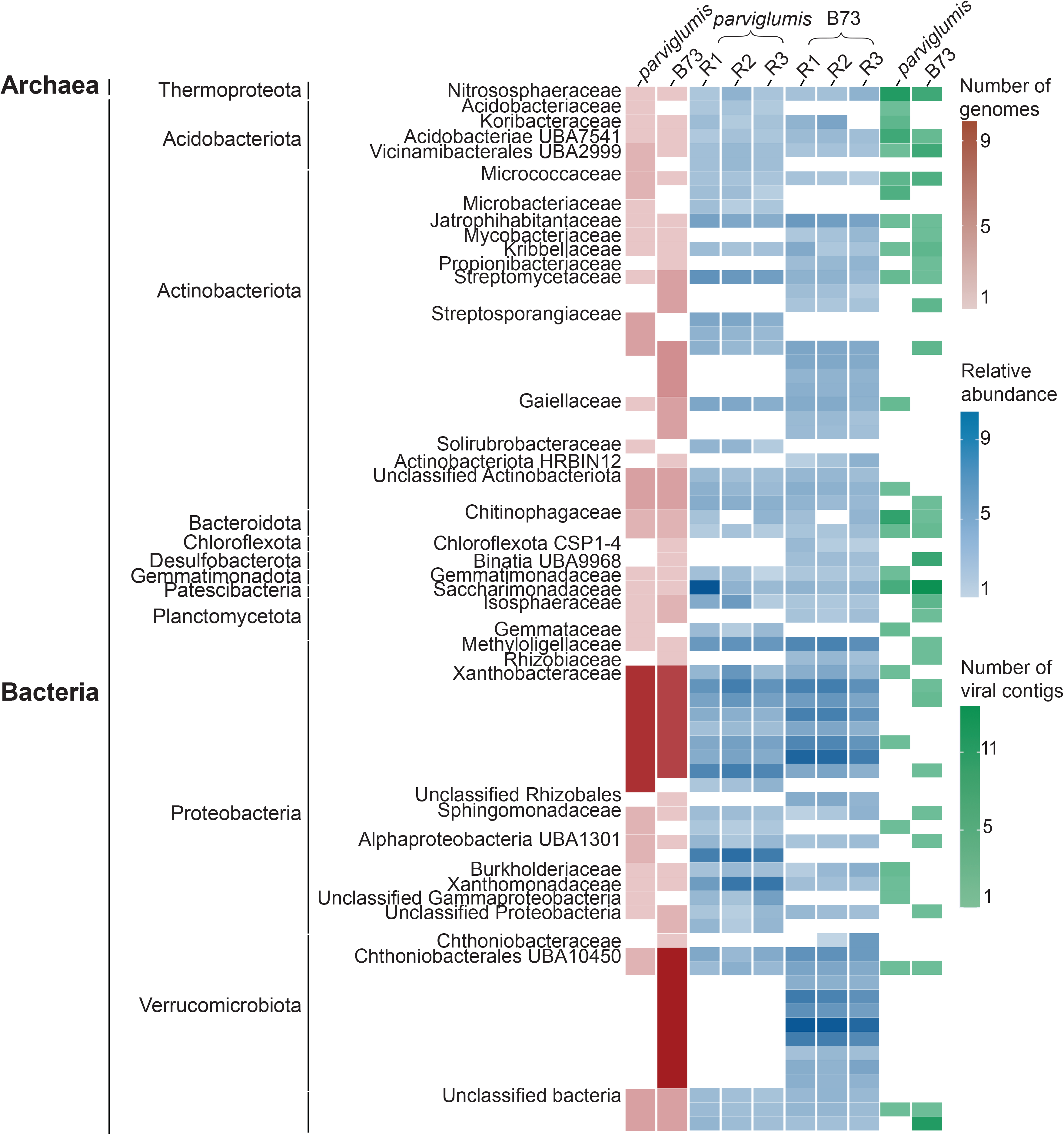
Metagenome assembled genomes (MAGs) and viral contigs from the Z. mays *parviglumis* and B73 rhizobiome. MAGs were reconstructed and viral contigs were identified from three biological replicates from each genotype, *parviglumis* and B73 rhizobiome. The number of genomes corresponding to the archaeal and bacterial taxa classified by the GTDB database and clustered to the family level when possible is denoted in red and the relative abundance of genomes is denoted in blue. The mean coverage depth of MAGs normalized against the number of reads in the sample to calculate the relative abundance of the MAGs.

As observed in 16S rRNA gene sequence data, taxa identified from MAGS revealed variance in the microbial community structure between genotypes (Fig. 2d; *p =* 0.001*, R^2^=* 0.99). The PCoA revealed 88.1% and 2.3% of the variance on axis 1 and 2, respectively (Fig. 2d). High variability of PCoA axis 1 demonstrated the variation in beta diversity between the two genotypes. This further highlighted the differences in the microbial community structure of *Z. mays* genotypes.

Similar to the 16S rRNA gene amplicon sequence data, MAGs also identified Proteobacteria and Actinobacteriota as the two most abundant rhizobiome phyla of both *parviglumis* and B73 rhizobiome (Fig. 3). Analyses of the B73 rhizobiome 16S rRNA gene amplicon data revealed that in addition to taxa identified in the Proteobacteria and Actinobacteriota, Verrucomicrobiota taxa were abundant community members (Fig. 2a, 2c, and 3). Within the Proteobacteria, community members in the family Xanthobacteraceae (18.00%) were the most abundant in the *parviglumis* rhizobiome. This result is different relative to the most abundant taxa identified at the family level in the B73 rhizobiome, where taxa within the Chthoniobacterales UBA10450 in the Verrucomicrobiota (16.39%) were the most abundant in B73 followed by Xanthobacteraceae (13.11%), Chloroflexota CSP1-4 (1.62%) and Binatia UBA 9968 in Desulfobacterota (1.63%) (Fig. 2c and 3). In addition to microbial taxa, viral contigs were also identified from the shotgun metagenome sequence data (Fig. 3).

### Rhizobiome viral community structure variation between parviglumis and B73

Identification of putative viruses from rhizobiome shotgun metagenome sequence data detected 329 and 488 viral contigs in *parviglumis* and B73, respectively. Further screening of viral contigs yielded 67 and 86 contigs with in the rhizobiome of *parviglumis* and B73, respectively. The average nucleotide identity (ANI) distribution not only displayed a high level of strain heterogeneity but also revealed a wide range of viral genome variations. The genome size of putative viruses identified in the parviglumis rhizobiome ranged from 1.50 – 40.54 Kbp with 5 vMAGs and B73 viral contigs ranged from 1.51 – 31.18 Kbp with only 1 vMAG reconstructed. Viral contig clustering between two rhizobiomes resulted in the identification of only 7 (4.5%) viral contigs identified in both *parviglumis* and B73 (2 clustered > 3 kb and 5 clustered < 3kb) (Fig. 4a). Rhizobiome viral community structure collected from each genotype was significantly different (*R^2^ =* 0.999*, p =*0.005) (Fig. 4b).

**Fig. 4.**
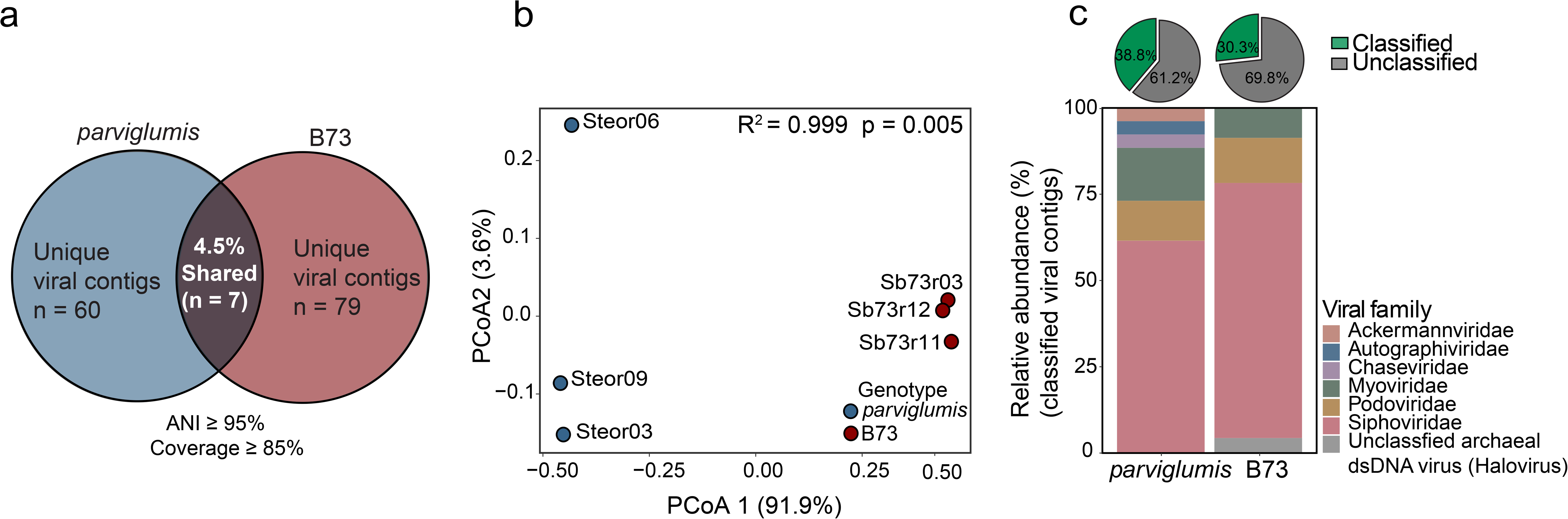
Identification of viral contigs within the *parviglumis* and B73 rhizobiome. To assess the similarities/differences of viral contigs in the *parviglumis* and B73 rhizobiome, pairwise comparison with average nucleotide identity (ANI) > 95% with coverage > 85% of the viral contigs of the shorter sequence was performed where (a) only 7 viral contigs were clustered > 95% ANI showing heterogeneity in identified viral contigs. (b) Ordination conducted with Bray-Curtis dissimilarity on the relative abundance of the viral contigs explained 91.9% variance on PCoA1 and 3.6% variance of PCoA2 (*R^2^* = 0.999 *p* = 0.005) as analyzed by permutational multivariate analysis of variance (perMANOVA) conducted utilizing adonis2. (c) Taxonomic classification of identified viral contigs comprised 6 different families of Caudovirales in *parviglumis* and 3 families in B73 with an unclassified archaeal virus. The relative abundance of the classified viral contigs was calculated as the proportion of the corresponding viral family divided by the total classified viral contigs multiplied by 100.

Taxonomy of rhizobiome viral contigs could be assigned to 38.80% in *parviglumis* and 30.23% in B73 viral contigs. Putative viruses were taxonomically classified as Caudovirales and among the annotated viral contigs, the family *Siphoviridae* (61.53%), *Myoviridae* (15.38%), and *Podoviridae* (11.53%) were the most abundant in the viral community of the *parviglumis* rhizobiome. The *Siphoviridae* (76.92%), *Podoviridae* (11.53%), and *Myoviridae* (7.69%) were prevalent in the B73 rhizobiome. Contigs classified in the families of *Ackermannviridae* (3.84%), *Autographviridae* (3.84%), and *Chaseviridae* (3.84%) were unique to *parviglumis* (Fig. 4c). One unclassified archaeal dsDNA virus, Halovirus (3.84%) was identified in the rhizobiome of B73 (Fig. 4c) but was not identified in *parviglumis* rhizobiome. Over two-thirds of the viral contigs identified remained unclassified and could not be assigned to a specific viral taxa.

### Virus-host linkages

Virus-host linkages could be established for 31.34% of *parviglumis* and 36.04% of B73 viral contigs. Phyla Thermoproteota, Acidobacteriota, Actinobacteriota, and Patescibacteria were commonly identified in both genotypes and were associated with the highest number of viral contigs in *parviglumis* and B73 rhizobiome. Few viral contigs were coarsely associated with unclassified bacteria (4.76% in *parviglumis* and 9.67% in B73). Thermoproteota family Nitrososphaeraceae was the only archaeal host identified in *Z. mays* rhizobiome. Of the identified hosts, 19.04% and 9.67% of viral contigs were linked to Nitrososphaeraceae in *parviglumis* and B73, respectively (Fig. 5a). Among the bacterial taxa Vicinamibacterales UBA2999, Micrococcaceae, Saccharimonadaceae, Xanthobacteraceae, and some unclassified bacteria were identified as common hosts in *parviglumis* and B73 (Fig. 5a). Viruses linked to Acidobacteriae UBA2999, Koribacteraceae, and Streptomycetaceae were unique to *parviglumis.* Rhizobiaceae, Propionibacteriaceae, Binatia UBA9968, Alphaproteobacteria UBA1301, Sphingomonadaceae, Chthoniobacterales UBA10450 were unique hosts in the rhizobiome of B73. None of the viral contigs were found to be infecting more than a single MAG in an established virus-host correlation indicating the virus host specificity at the MAG level. Although Verrucomicrobiota member Chthoniobacterales UBA10450 was highly abundant in the rhizobiome of both genotypes, it served as a virus host only in the B73 rhizobiome. Together these results indicate the variation of the community in the rhizobiome with host specificity.

**Fig. 5.**
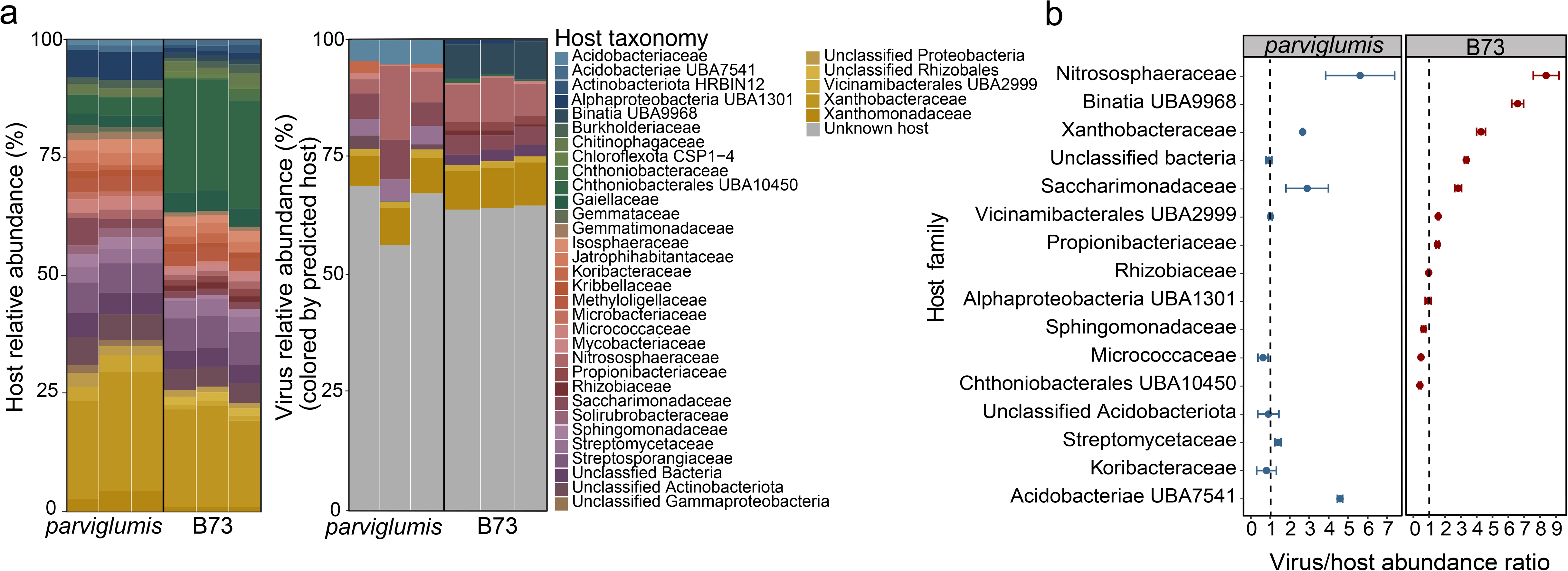
Virus-host linkages identified in the *parviglumis* and B73 rhizobiome. (a) Viral contigs were linked to the hosts utilizing genome similarities, CRISPR sequences, and tetranucleotide frequencies between the metagenome-assembled genomes (MAGs) and viral contigs. While the hosts identified for the *parviglumis* and B73 viral contigs showed differences in the host preferences, hosts couldn’t be identified for more than 60% of viral contigs in both genotypes and are identified in gray. (b) Virus to host abundance ratio was divided into 3 categories; less than 1, approximately 1, and greater than 1 demonstrated variation in viral genome copy number to their host in *parviglumis* and B73.The black vertical line separates the *Z. mays* genotypes, *parviglumis* and B73. The viral contigs are color-coded according to the identified potential host. The host taxonomy is identified at the family level (or lowest level of classification). Error bars represent the standard error of measure.

A total of 64 and 98 putative CRISPR spacer regions were detected in *parviglumis* and B73 infected MAGs, respectively. Among all the identified CRISPR spacer sequences in both genotypes, only a single CRISPR spacer sequence provided a positive hit to one of B73 MAGs. This corresponded to B73 MAG Xanthobacteraceae VAZQ01 sp005883115 with a 100% nucleotide identity match. The direct repeats flanking the CRISPR spacer region compared against the infected MAGs rendered an e-value < 10^-10^ confirming infection in Xanthobacteraceae VAZQ01 sp005883115 in B73 rhizobiome.

The average virus to host abundance ratio ranged from 0.41 to 8.38 in rhizobiome *parviglumis* and B73. Viruses infecting the Micrococcaceae (Actinobacteriota) were lower in abundance than hosts in rhizobiome collected from both genotypes. Viruses infecting Koribacteraceae (Acidobacteriota) in *parviglumis* and Chthoniobacterales UBA10450 (Verrucomicrobiota) in B73 were also lower than host cell. Virus abundance equaled host cell abundance of Vicinamibacterales UBA2999 (Acidobacteriota) in *parviglumis* only. Rhizobiome viruses exceeded the host cells, Nitrososphaeraceae (Thermoproteota), Xanthobacteraceae (Proteobacteria), and Saccharimonadaceae (Patescibacteria) in both genotypes(Fig. 5b).

### Metabolic potential of microbial and viral rhizobiome

The metabolic potential of the microbial community revealed the ability of the community to degrade carbohydrates as well as short chain fatty acids (Fig. 6). Bacterial hosts such as Xanthobacteraceae (Proteobacteria) not only showed the capability to degrade polyphenols, starch, short chain fatty acids, alcohols, chitins, and acetate utilization but were also capable of central carbon, nitrogen, sulfur, and iron metabolism (Fig. 6). Host cells Acidobacteriota (Acidobacteriae UBA7541, Vicinamibacterales UBA2999, Koribacteraceae) and Actinobacteriota (Micrococcaceae, Propionibacteriaceae) harbored genes that associated with polysaccharide degradation enabling central carbon metabolisms such as glycolysis and the Kreb’s cycle. Interestingly, the metabolic potential for CO_2_ fixation, utilizing the dicarboxylate-hydroxybutyrate cycle, hydroxypropionate-hydroxybutylate cycle, and reductive citrate cycle (Arnon-Buchanan cycle) were identified in MAGs corresponding to archaea Thermoproteota and bacteria Actinobacteriota, Bacteriodota, Proteobacteria, Verrucomicrobiota in both genotypes. Genes associated with the Calvin Cycle were not identified in any of the MAGs. The metabolic potential to degrade polyphenol and cleavage of arabinan and fucose cleavage was restricted to the Koribacteraceae identified in the *parviglumis* rhizobiome (Fig. 6a). The ability to degrade chitin was specific to Desulfobacterota (Binatia UBA9968) and Verrucomicrobiota (Chthoniobacterales UBA10450) with the potential to degrade beta-mannan in the B73 rhizobiome (Fig. 6b). Archaea Thermoproteota indicated the metabolic potential to convert acetate into methane, carbohydrate degradation and detoxification of aromatic compounds, and ammonia oxidation. Additionally, *parviglumis* MAGs Thermoproteota (Nitrososphaeraceae) and Proteobacteria (Xanthobacteraceae) were capable of denitrification. No B73 MAGs showed any role in nitrification and similarly, no *parviglumis* or B73 viral infected MAGs showed any role in sulfur metabolism. The microbial community in both the *parviglumis* and B73 viral infected MAGs showed a role in siderophore transport to support iron acquisition in soils.

**Fig. 6.**
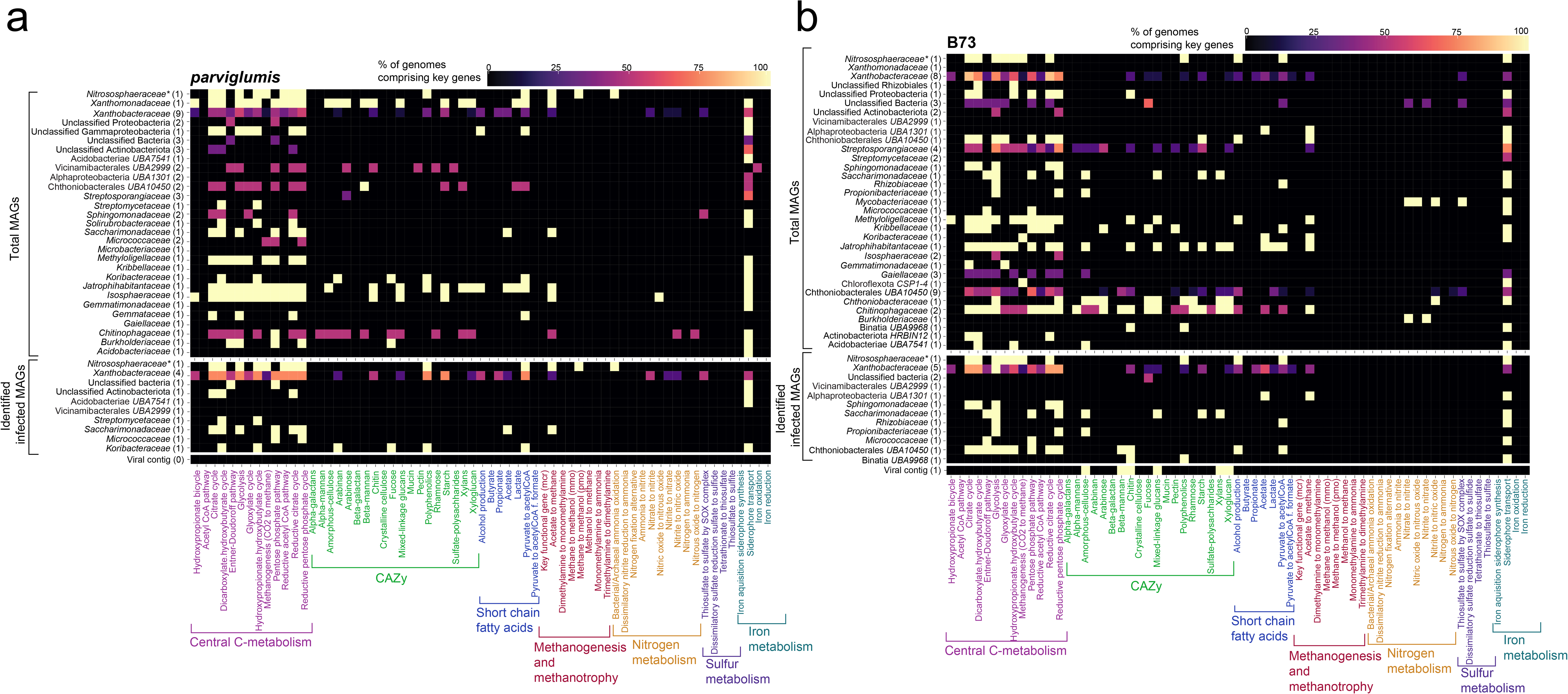
Metabolic potential of the microbial community identified from the annotation of MAGs and virus contigs in the *parviglumis* and B73 rhizobiome. The rhizobiome metabolic potential was determined using DRAM, METABOLIC, and FeGenie. The heatmap represents the metabolic potential of the total metagenome-assembled genomes (MAGs) (top), viral infected MAGs (middle), and contribution of viral auxiliary metabolic genes (AMGs) (bottom) of (a) *parviglumis* and (b) B73 rhizobiome, respectively. Percentage of genomes were determined at the family level.

One auxiliary metabolic gene was identified in a viral contig collected from the rhizobiome of B73 whereas AMGs were not identified in viral contigs recovered from *parviglumis*. Analysis of the viral encoded AMG identified this gene as a glycoside hydrolase family 5 (GH5) capable of degrading amorphous cellulose, xyloglucan, xylan, beta-mannan, mixed-linkage glucans backbone cleavage, and chitin (Fig. 6b). This virus was linked to the host Propionibacteriaceae (Actinobacteriota) in the B73 rhizobiome. These results indicate that rhizobiome viruses can harbor genes with the potential to produce labile mono and oligosaccharides.

## Discussion

Here we demonstrated that both microbial and viral community structure varies between two *Z. mays* genotypes, *parviglumis* and B73, cultivated in the same homogenized soil under the same environmental conditions. While it is recognized the *Z. mays* genotype influences the rhizosphere microbial community [71], to our knowledge this is the first study to demonstrate the variation of the rhizobiome viral community structure between plant genotypes. Plants release root exudates below ground, creating a diverse chemical milieu that impacts rhizobiome assemblage and structure [46, 72, 73]. The root exudate composition is also recognized to vary among maize genotypes [25], thus further promoting variation in the rhizosphere community. Our results are consistent with prior research demonstrating *Z. mays* genotype plays a factor influencing the selection of rhizobiome microbial community [71]. Here we identified Proteobacteria, Actinobacteriota, and Verrucomicrobiota constituted the core microbiome of *parviglumis* and B73 consistent with a prior study [46] and were more abundant relative to the Acidobacteriota and Desulobacterota. Interestingly, Acidobacteriota were associated with more viral contigs which suggests an active infection could result in lower cell abundance. The relatively lower abundant microbial taxa in *Z. mays* rhizobiome such as Chloroflexota and Desulfobacterota (relative abundance < 2.5% in 16S rRNA gene data) were reconstructed in B73, but we were unable to recover MAGs from *parviglumis*. This could be a result of low relative abundance and/or lysed host cells not detected in sequencing. Alternatively the result could also be a result of differential root-exudate production between genotypes [25] and recruitment of these taxa to the root. Microbial community structure and viral host cells are recognized to influence viral community structure [31, 74, 75]. Small but significant differences in rhizobiome microbial community composition in *parviglumis* and B73 manifested dramatic differences in the viral community of these two *Z. mays* genotypes.

Viruses were abundant in the rhizobiome of *parviglumis* and B73 rhizobiome outnumbering cells. The virus to cell ratio resulted in a VCR >1.5, 3 times more than previously reported in the wheat rhizosphere (VCR = 0.27) [8]. Viral contigs recovered from the shotgun metagenome sequence data supported variation of the viral community between genotypes revealing the presence of a distinct viral community in the rhizobiome of *parviglumis* and B73. We recognize that members of the viral community composition reported in this study may represent an underestimate of viral diversity in the *Z. mays* rhizobiome due to sampling constraints because reported data in this study are limited to viruses associated with the host genome, adsorbed to particulate matter, and greater than 0.2 μm in size. Prior research has demonstrated significant variations in both microbial and viral community due to spatial distribution and/or soil heterogeneity [59, 76, 77], the close clustering of the *Z*. *mays* biological replicates on PCoA plot demonstrated that soil heterogeneity among replicates didn’t contribute to significant differences in microbial and viral community structure. Rather the genotype of the plant played a significant role in the pronounced variation in the rhizobiome viral community where only seven viral contigs were shared between the two plant genotypes. This finding is also supported by pairwise comparison clustering of viral contigs which yielded only 7 similar viral contigs and validated that *parviglumis* and B73 comprised contrastingly different viral communities while grown in the same conditions using the same soil inoculum. Thus in addition to soil physiochemical factors, plant species and genotype can further contribute to the spatial variation of viral communities observed in soils.

### Viral community variation in parviglumis and B73

Virus-host specificity and host range abundance play an important role in viral selection in the rhizosphere [8]. The viral contigs recovered from the *parviglumis* and B73 rhizobiome revealed host specificity. All viral contigs recovered were found infect a unique MAG. The strong correlation between cell and virus abundance is interconnected through prey-predator interactions. Repeated viral infections and host cell lysis give rise to virus-defense through immunity systems including the CRISPR-cas system found within some bacterial and archaeal lineages, thus, avoiding future viral infections [78]. The presence and 100% identity match to the CRISPR sequence in B73 MAG suggested the adaptation of a defense mechanism against viral infection by Xanthobacteraceae. This active defense mechanism of bacteria exploiting CRISPR sequences could be a possible explanation for the lower number of viral contigs in Xanthobacteraceae MAGs hence, the high abundance of Xanthobacteraceae in rhizobiome. The small genome size of the ultra-small bacteria Saccharimonadaceae (Patescibacteria) may not be able to comprise CRISPR sequences thus rendering the cell more susceptible to viral infection [79, 80]. However, the Patescibacteria adopted the strategy to evade viral infection by deleting common membrane phage protein receptors [81]. Additionally, the choice of viral reproduction strategy between lytic and lysogeny can affect the host community dynamics resulting in distinct fates of host cells [4, 82, 83]. Viral host specificity can thus play a role influencing microbial diversity [75]. Viral predation results in altered community structure [84], can increase the release biologically available nutrients, and can consequently impact ecosystem productivity [85]. Here we observed rhizobiome viral contigs linked to the host cells which demonstrated host preference which could subsequently impact host abundance as demonstrated by the variation in the ration of viruses abundance to host abundance. Viral infection found to be impacting host metabolic output not only by impacting host abundance but also through acquiring glycoside hydrolases AMG capable of complex polysaccharide degradation [15, 86].

### Impact of virus-host interactions on metabolic potential

Viruses that exhibit a lytic life cycle cause cell-mediated cell lysis liberating labile sources of carbon and nutrients [87] which can stimulate microbial population growth from the bottom up potentially altering community diversity and metabolic output [88, 89]. In a stable isotope probing (SIP) study performed in soils, the phyla Acidobacteriota, Actinobacteriota, Chloroflexota, Proteobacteria, Patescibacteria, Verrucomicrobiota were identified as active viral hosts involved in soil carbon cycling [80]. Taxa identified in our study were also identified to harbor the metabolic potential to drive carbon cycling near plant roots. Acidobacteriota (Acidobacteriae UBA7541, Vicinamibacterales UBA2999, Koribacteraceae), Actinobacteriota (Micrococcaceae, Propionibacteriaceae, Streptomycetaceae), Patescibacteria (Saccharimonadaceae), Proteobacteria (Alphaproteobacteria UBA1301, Sphingomonadaceae, Rhizobiaceae, Xanthobacteraceae), Verrucomicrbiota (Chthoniobacterales UBA10450), with the addition archaeon Thermoproteota (Nitrososphaeraceae) are involved in central carbon metabolism. In addition to carbon biogeochemical cycling, members of the rhizobiome of both *parviglumis* and B73 had the potential for siderophore transport to support iron acquisition in soils. We did not detect the metabolic potential for nitrification and sulfur metabolism within taxa that were infected by viruses.

Viral encoded glycoside hydrolases AMGs have the potential to contribute to carbon metabolism in *Z. mays* rhizobiome through expression within the host cells. Here we observed the unclassified vMAG in the B73 rhizosphere that contained a glycoside hydrolase (GH5) which is recognized to degrade amorphous cellulose, beta-mannan, chitin, mixed-linkage glucans, xylans, and xyloglucan. The acquisition of this AMG, glycoside hydrolases responsible for carbohydrate hydrolysis, by soil viruses is consistent with prior studies [15, 35, 80, 90] and demonstrates the potential within the rhizobiome. This vMAG that was linked to host Propionibacteriaceae (Actinobacteriota) that respond to a variety of carbon sources by secreting extracellular enzymes such as cellulases, amylases, and chitinases were found to be regulating carbohydrate catabolic pathways [91, 92]. These results indicate that virus-encoded glycoside hydrolases have the potential to contribute to the production of labile mono and oligosaccharides which can further serve as a carbon source for the host and other rhizobiome community members.

## Conclusion

Together these results indicated that plant genotype plays a role in microbial assemblage in rhizobiome and viral community parallelly shifts in response to microbial community selection. Viral community structure is linked to compositional patterns of microbial communities based on plant genotypes. Thus, variation in below ground plant roots has the potential significantly influence heterogeneity of viruses within soils. The virus-host interaction showed differences in host abundance between two genotypes which has the capability to influence the metabolic potential of the rhizobiome. Viral AMGs are capable of carbohydrate degradation which have the potential to produce labile mono and oligosaccharides in the rhizosphere. These labile carbon sources could further promote microbial community proliferation. Therefore, to extent to which host-virus interaction alters host abundance, microbial community structure, and functions remains a critical question in rhizosphere microbial ecology and soil biogeochemical cycling.

## Supporting information

Experimental Data

## Acknowledgements

This project was funded by the National Science Foundation EPSCoR Center for Root and Rhizobiome Innovation Award #1557417 to KAW and SER.

## Data availability

Experimental data and results summary have been uploaded as a supplement. DNA sequence data and corresponding metadata has been deposited to the NCBI under BioProject accession number PRJNA955173.

## Author’s contribution

PY and KAW designed the study. PY, AQ, YY, TK, SER, and KAW developed the data collection protocols. PY, AQ, YY, TK, JO, RK, SER, and KAW conducted the experiment and collected the data. PY and KAW analysed the data, with help from SER and AQ, PY and KAW wrote the initial draft. All authors edited the manuscript.

