## Supplementary Information for "Zea mays genotype influences microbial and viral rhizobiome community structure"

**Methods**

*Seed germination and soil*

*Z. mays spp. parviglumis* is an ancestral maize lineage commonly known as teosinte whereas *Z. mays* genotype B73 is an early maize hybrid. Seeds were surface sterilized using 2% TWEEN (polysorbate) solution for 90 seconds followed by priming incubation in aerated sterile 1 mM calcium chloride solution for 24 hours. The seeds were germinated in sterile petriplates lined with autoclaved Kimwipes soaked in 1 mM calcium chloride solution in a dark growth chamber at 25° C [1]. Once seeds germinated, they were transplanted into the experimental soil matrix.

Soil was collected from the top 30 cm of a native tallgrass prarie, Nile Mile Prairie, sieved (2 mm mesh), homogenized, and mixed with autoclaved sandy agricultural soil also collected from the top 30 cm from the UNL Agricultural Research and Development Center (ARDC) in Mead, Nebraska (41°09'31.3"N 96°24'47.4" W), sieved (2 mm mesh), and homogenized. The agricultural sandy soil was autoclaved for two dry cycles of 121°C and 110 kPa for 100 minutes and was kept covered in autoclaved sterile aluminum containers (< 45 Kg per container) to avoid environmental contaminants until mixed with live prarie soil. The soil mixutre was allowed to equilibrate in sterile sealed aluminum containers for a week prior to potting.

*Below ground plant phenotype data collection*

Root phenotype parameters were collected using WinRhizo root scanning software within one week of plant harvesting. In the event any soil remianed on the root surface, soil was gently removed and roots were untangled using forceps to avoid overlap of the root hairs without breaking the roots. Roots were stained in Toluidine blue (0.05% in pH 4.4) and positioned in the WinRhizo Epson Perfection scanning tray immersed in double distilled water with minimum overlaps and illuminated from the top and bottom sides to scan the roots. The scanned root images were used to evaluate root phenotype parameters such as length, diameter, and volume using Winrhizo image analysis software (Regent Instruments, version 2008a). To differentiate the allocation traits precisely, roots were categorized into coarse (> 1 mm) and fine (≤ 1 mm) roots in the WinRhizo root analysis program. For the roots, based on the root category, coarse and fine roots fresh weight (g), total below ground fresh and dry weight (g), root dry matter content coarse and fine roots (g/g), coarse and fine root length (cm), total root length (cm), the surface area of coarse and fine roots ($cm^2$), total root surface area ($cm^2$), coarse and fine root volume ($cm^3$), total root volume ($cm^3$), total biomass in dry weight (g) was collected.

Intrinsic root traits such as specific root length for coarse and fine roots (cm/g), coarse and fine root density ($g/cm^3$), total root density ($g/cm^3$), average coarse and fine root diameter (cm) were calculated as previously described [2]. Plant growth rate data comprise growth coarse and fine root growth rate (cm/day), and total root length growth rate (cm/day). Allocation traits such as coarse and fine root length ratio (cm/g), root mass ratio (g/g), total length ratio (coarse and fine roots), below-ground plant ratio (g/g), below-ground mass ratio (g/g) was calculated.

*Cell and virus enumeration*

Cell and virus enumeration from biological replicates of *parviglumis* (n = 5) and B73 (n=6) were collected. One of the *parviglumis* biological replicates was discarded as a result of an error occurring during sample collection. Following sample collection, all cell and virus samples were enumerated within one week. Cell samples were collected on an Isopore 0.2 µm black polycarbonate filters (product# GTBP02500), while viruses were collected on 0.02 µm Anodisc™ filters (product# 6809-6002) for enumeration with Whatman 0.45 µm cellulose nitrate membrane filters (product# 7184-002) as a supported filter to ensure uniform distribution of cells and viruses. Samples were passed through each filter by using Gast high-capacity vacuum pump (model# DOA-P704-A4) maintaining a constant pressure of less than -0.2 bar to avoid cell lysis. SYBR Green I was used to stain the filters for 15 minutes followed by drying [3]. The filters were mounted on slides with an anti-fade solution (50% glycerol, 50% PBS, 0.1 p-phenylenediamine). At least 200 particles in 10 field views were enumerated per filter [4] and averaged to estimate the cells and viruses in the rhizobiome. Later the enumeration data was analyzed against rhizosphere mass and other mentioed root parameters.

*Nucleic acid normalization for 16S rRNA gene amplicon and shotgun metagenomic sequence library preparation*

All the glassware used to extract rhizobiome (rhizosphere and rhizoplane collectively) was treated with 0.1% diethylpyrocarbonate (DEPC) whereas spatulas, needles, and other equipment such as sonicator horn dip were sterilized by treating them with bleach (10% v/v), ethanol (70% v/v), distilled water (x3), RNase AWAY™ surface decontaminant (product#7002), and finally with 0.1% DEPC treated water (x3) rinse.

$$DNA\ per\ gram\ of\ rhizobiome\ \left(\frac{ng}{g}\right) = Total\ DNA\ extract\ (uL) \times$$

$$DNA\ concentration\ \left(\frac{ng}{uL}\right) \div\ Rhizbiome\ mass\ (g)\ used\ for\ DNA\ extraction$$

*Bioinformatic analyses: 16S rRNA gene amplicon sequences*

The 16S rRNA gene amplicon sequence data was analyzed using the DADA2 package [5] with parameters (truncLen=c(240,160), maxN=0, maxEE=c(2,2), truncQ=2, rm.phix=TRUE) to obtain amplicon sequence variants (ASVs). ZymoBiomics microbial community standard (product # D6300) with known microbial community and abundance served as a positive control for 16S rRNA gene amplicon sequencingw. The mentioned parameters for the 16S rRNA amplicon sequencing processing pipeline were validated using a positive control to detect contamination and remove spurious sequences (ASV abundance < 0.05 %). The diversity and evenness were measured using Shannon's diversity index and Pielou's evenness, respectively, and observed richness was the total number of ASVs. Broad SILVA v138 (Silva 138 SSURef Nr99) [6] was used to assign taxonomic identification to the ASVs.

*Bioinformatic analyses: Shotgun metagenome sequences*

The metagenomic raw reads were quality checked to exploit FastQC and Trimmomatic v0.39 [7] was used to remove Illumina adapter sequences and quality trim with parameters (LEADING:2 TRAILING:2 SLIDINGWINDOW:4:20 MINLEN:25 ILLUMINACLIP:NexteraPE-PE.fa:2:40:15) and PhiX sequences were removed using BBDuk v38.84 [8]. To ensure the quality of the clean reads FastQC check was performed again. The clean reads were used for three biological replicates of the *parviglumis* and B73 utilizing MEGAHIT v1.2 [9], using a contig minimum size threshold of 1000bp. Bowtie2 v2.4 [10] was used to map the reads back to the assembly and indexed using SamTools v1.9. [11]. The final set of MAGs were categorized based on Minimum Information about a Metagenome Assembled Genome (MIMAG) quality parameters [12]. The relative abundance of the reconstructed MAGs was estimated by normalizing the mean coverage depth of metagenome reads against the total reads of the samples [13-15]. Taxonomic identification of reconstructed MAGs was performed using 22 single copy core genes coding for ribosomal proteins using Genome Taxonomy Database (GTDB) [16-19].

*Viral contig detection and identification*

To identify the viral contigs VirSorter 2 (v2.1) was used (--high-confidence-only --hallmark-required --prep-for-dramv) [20]. To retrieve only the genuine viruses output of VirSorter2 was facilitated as input of DeepVirFinder which identifies viruses based on reference-free and alignment-free machine learning methods [21]. The viral contigs identified by DeepVirFinder comprising p- value <0.05 were retained. SortmeRNA [22] was used to detect the 16S rRNA fragment contamination in the viral contigs. The final set of viral contigs was mapped using Bowtie2 v2.4 [10] and resulting alignments were sorted and indexed using SamTools v1.9. [11]. BamM 'parse' coverage [23] with 'tpmean' coverage mode was used to generate abundances of the MAGs. The putative viral contigs Kaiju (v1.8) [24] and BLASTN (e-value $\leq 10^{-5}$, bit score $\geq$ 50) against NCBI-viral RefSeq was used to taxonomically identify viral contigs. Putative viral contig sequences queried against the NCBI ref database with BLAST [25] with threshold e-value $< 10^{-5}$ was also used to remove any bacterial contamination.

*Virus-host linkage*

To validate the viral infection of a host from sequence identified in MAGs, sequences were detected in the MAGs utilizing CRISPRCasFinder [26]. The retrieved spacers were matched against all viral contigs using BLASTN and matches with 0 mismatches and e-value $\leq 10^{-5}$ hit were scored as a positive match. For the spacer with the positive hit, the direct repeat sequences were compared to the MAGs via BLASTN threshold e-value $10^{-10}$ and 100% nucleotide identity to link between the CRISPR region and the host [27]. The BLASTN (e-value $10^{-5}$, bit score $\geq 50$) was used to link viral contigs to the hosts, based on their shared genomic contents such as AMGs and/or tRNAs [28, 29]. Normalized virus-host relative abundance was calculated by dividing viral contig relative abundance by viral-linked host abundance at the family level (or at order/class/phylum level if the family was not informative) [15, 27].

### Supplementary Figures

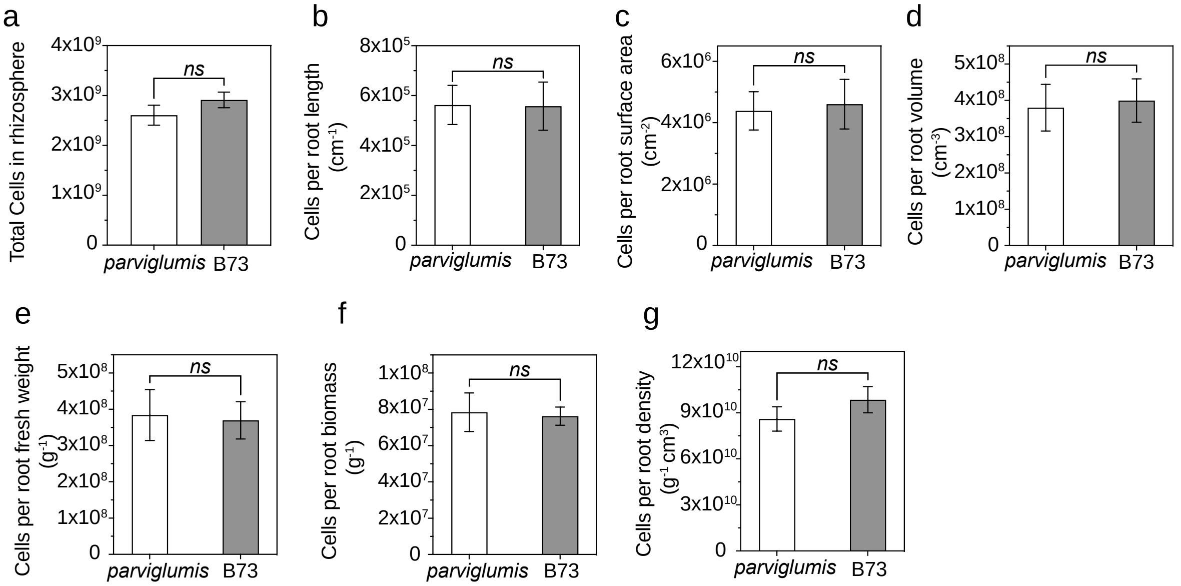


**Fig. S1 Cell enumeration data against other root parameters.** Statistical differences were quantified in the cell abundance of biological replicates of *parviglumis* (n=5) (transparent) and B73 (n=6) (gray) enumerated using epifluorescence microscopy with SYBR Green I dye. The (a) represents the total cells collected in rhizosphere and cell enumeration against (b) root length, (c) root surface area, (d) root volume, (e) root fresh weight, (f) root biomass, and (g) root density. The non-parametric unpaired t-test with Welch's correction was conducted assuming unequal variances did not show any significant difference between the *parviglumis* and B73 cell count data against any of the analyzed root parameters. Error bars represent standard error mean.

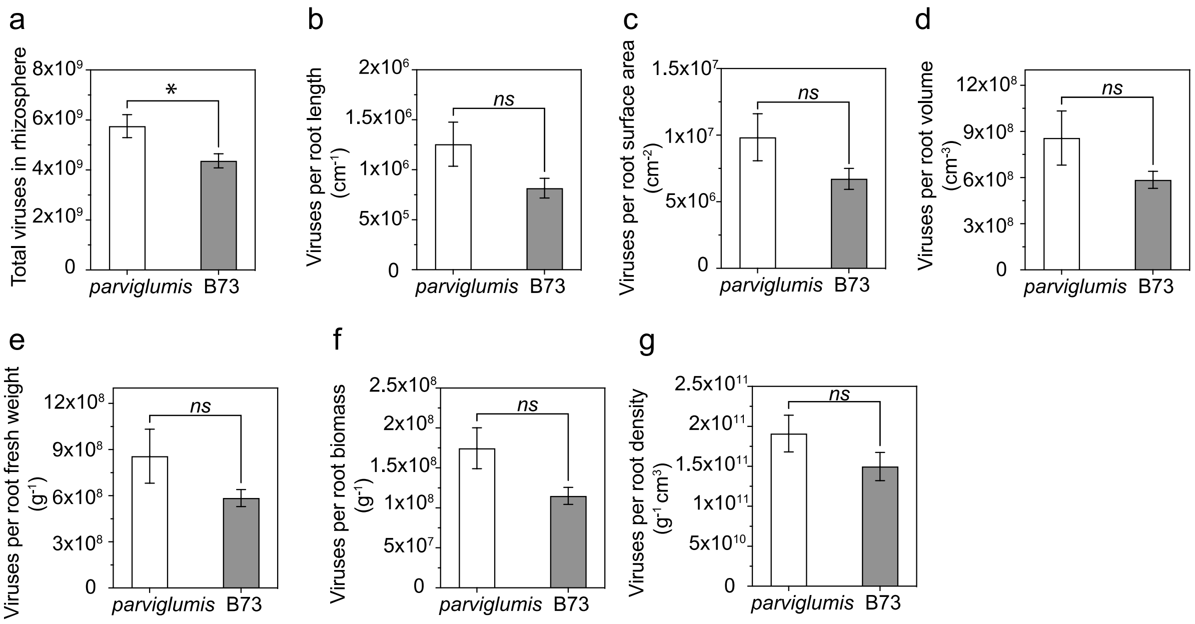


**Fig. S2 Virus enumeration data against other root parameters.** Statistical differences were quantified in virus abundance of biological replicates of *parviglumis* (n=5) (transparent) and B73 (n=6) (gray) enumerated using epifluorescence microscopy with SYBR Green I dye. The (a) represents the total viruses collected in rhizosphere and virus enumeration against (b) root length, (c) root surface area, (d) root volume, (e) root fresh weight, (f) root biomass, and (g) root density The non-parametric unpaired t-test with Welch's correction was conducted assuming unequal variances showed significant variation ($p = 0.0376$) while viruses against other root parameters did not show any significant difference between the *parviglumis* and B73 cell count data against any of the analyzed root parameters. Error bars represent standard error mean.

**Supplementary Information References**

[1] Rehman Ur H, Iqbal H, Basra SM, Afzal I, Farooq M, Wakeel A *et al*. Seed priming improves early seedling vigor, growth and productivity of spring maize. *J Integr Agr* 2015;14**:** 1745-1754.

[2] Quattrone A, Lopez-Guerrero M, Yadav P, Meier MA, Russo SE, Weber KA. Interactions between Root Hairs and the Soil Microbial Community Affect the Growth of Maize Seedlings. in review.

[8] Bushnell B (2014). BBTools software package.

[16] Starr EP, Shi S, Blazewicz SJ, Probst AJ, Herman DJ, Firestone MK *et al*. Stable isotope informed genome-resolved metagenomics reveals that Saccharibacteria utilize microbially-processed plant-derived carbon.

[17] Probst AJ, Ladd B, Jarett JK, Geller-McGrath DE, Sieber CMK, Emerson JB *et al*. Differential depth distribution of microbial function and putative symbionts through sediment-hosted aquifers in the deep terrestrial subsurface. *Nat Microbiol*. 2018;3**:** 328–336.

[18] Sharon I, Battchikova N, Aro EM, Giglione C, Meinnel T, Glaser F *et al*. Comparative metagenomics of microbial traits within oceanic viral communities. *ISME J*. 2011;5**:** 1178-1190.

[19] Parks DH, Chuvochina M, Rinke C, Mussig AJ, Chaumeil P-A, Hugenholtz P. GTDB: an ongoing census of bacterial and archaeal diversity through a phylogenetically consistent, rank normalized and complete genome-based taxonomy. *Nucleic Acids R*. 2022;50**:** D785-D794.
